# A Unified Dynamical-Systems and Control-Theoretic Model for Single-Cell Fate Dynamics

**DOI:** 10.64898/2026.03.15.711134

**Authors:** David M. Redd, Samuel G. Green, Tommy W. Terooatea

## Abstract

Single-cell technologies now resolve cell-fate transitions, yet most analyses remain descriptive rather than predictive. This model unifies pseudotime (geometry), RNA velocity (local direction), optimal transport (OT; distributional evolution) and Schrödinger-bridge approaches (stochastic trajectories) within a stochastic dynamical-systems and control lens under partial observability^1,2,3,4^.

We provide a minimal bridge from Chemical Master Equation models to SDE/Fokker-Planck descriptions and quasi-potentials, clarify what can and cannot be identified from snapshot data, and outline practical experimental design (time points, modalities, perturbations)^5,6,7^.

We summarize method-specific assumptions, strengths and failure modes; present case studies (iPSC reprogramming, pancreas endocrinogenesis, hematopoiesis at scale); and detail a 10-step workflow with uncertainty propagation and reporting standards^3,8,9^.

Finally, we recast intervention as a control problem: the realistic objective is probabilistic programmability-shifting terminal fate distributions with minimal inputs and preserved viability-rather than deterministic state-to-state command^10^.

## Introduction

Single-cell RNA-seq and multimodal assays (e.g., RNA+ATAC+protein) have made it routine to map cellular heterogeneity across development, regeneration and disease. However, snapshot sampling prevents direct observation of cell trajectories. This limitation catalyzed four influential families of analysis: pseudotime methods that reconstruct progression geometry; RNA velocity that estimates local temporal derivatives from splicing kinetics; optimal transport that couples successive population snapshots to infer probabilistic fates; and Schrödinger-bridge approaches that interpolate stochastically between marginals to separate drift from diffusion^1,2,3,4^.

This model connects these tools to a unified perspective: cells as stochastic control systems under partial observation, time courses as distributional evolution and perturbations (e.g., CRISPR, ligands, TF overexpression) as control inputs^5,6,10^.(Fig. 1)

**Figure 1.**
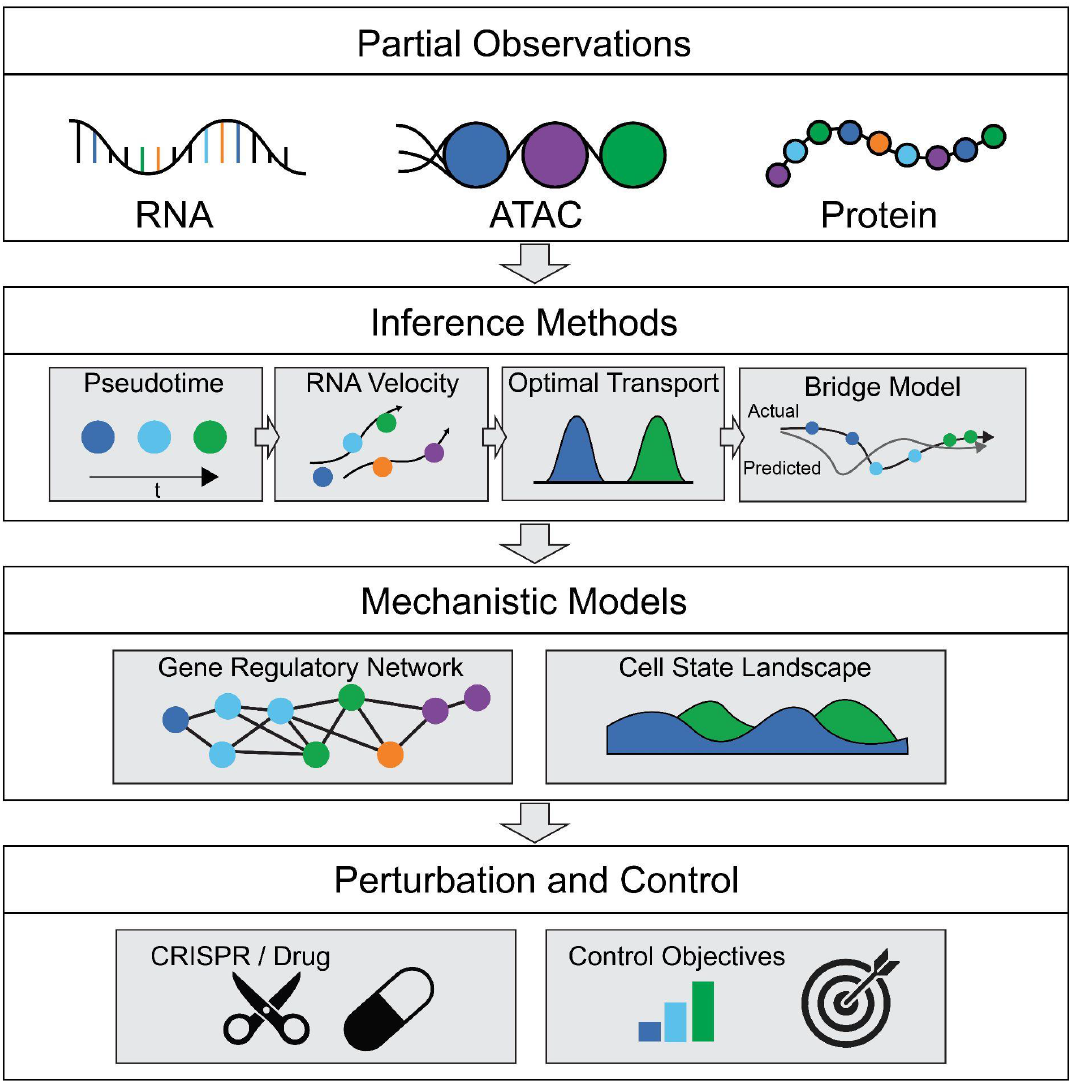
Four-layer schematic. Schematic linking measurement, inference, mechanism and intervention. RNA/ATAC/protein provide partial observations of latent state. Inference tools (pseudotime, velocity, OT, Schrödinger bridges) estimate geometry and distributional dynamics. Mechanistic interpretation invokes GRNs and landscape structure. Perturbations act as control inputs that reshape transition kernels and terminal fate distributions under partial observability.

## Experimentation

### Conceptual foundation and notation

We model cells as partially observed stochastic control systems: 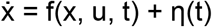, y = h(x) + ϵ, where x is latent cellular state, y are observations (RNA/ATAC/protein), u are perturbations, η captures biological noise and ϵ measures noise. In mesoscopic regimes, CME dynamics admit SDE approximations whose densities obey Fokker–Planck equations; under small noise with metastability, quasi-potentials approximate attractor basins and transition saddles; most biological systems are non-equilibrium and need not be globally gradient^5,6,7^.

Partial observability implies fundamental non-identifiability: multiple drifts/diffusions or GRNs can explain the same snapshots. Therefore, explicit assumptions (priors), orthogonal measurements (time labels, lineage, metabolic labeling) and perturbational constraints are essential.

### Data acquisition and study design

Time points: sample early, branching and terminal stages; add intermediate points near decision boundaries. If feasible, incorporate metabolic labeling to inform transcription/degradation rates and improve velocity calibration^2^.

Modalities: combine RNA with ATAC and/or protein to improve observability, anchor GRN hypotheses and adjudicate ambiguous transcriptional states.

Replication and controls: include replicates; record experimental time labels and lineage when possible; capture perturbation metadata.

Perturbations as inputs: design a hypothesis-driven panel (TFs, ligands, CRISPRa/i) to test predicted levers and to evaluate control objectives (increase desired lineage probability while maintaining viability)^10^. (Fig. 2)

**Figure 2.**
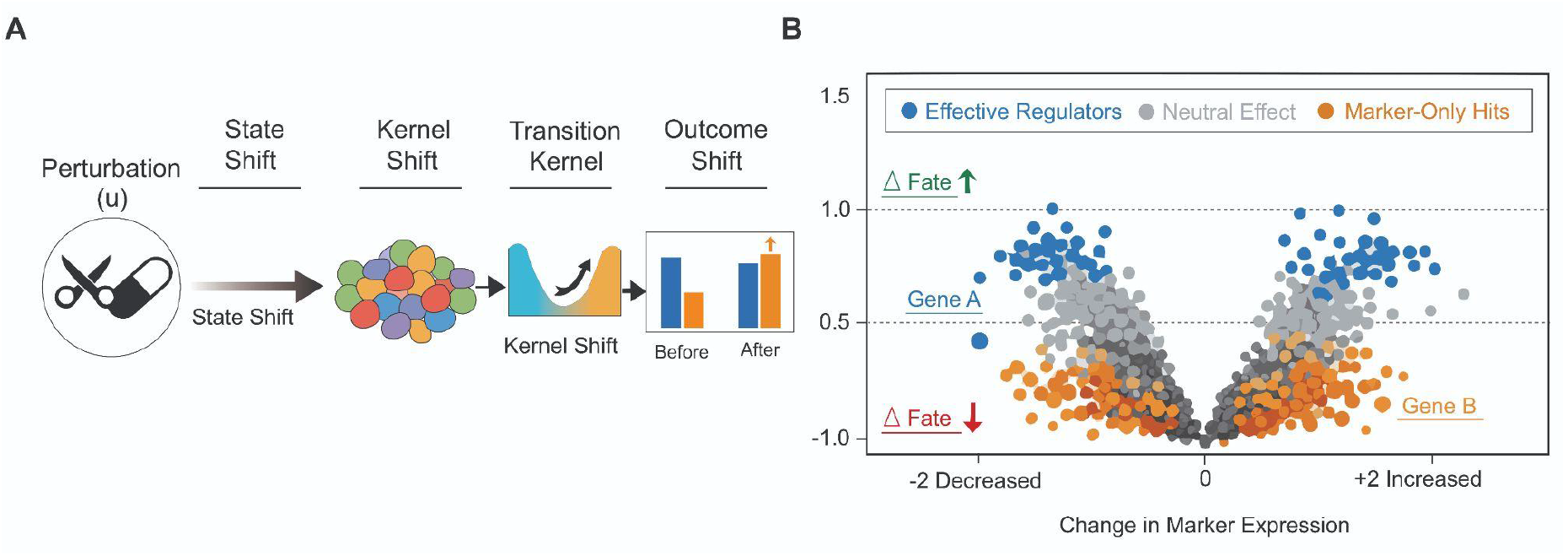
Perturbation as control input. (A) Schematic: perturbation input → state shift → transition kernel shift → altered fate probabilities. (B) Perturb-seq example showing perturbation-specific transcriptional programs and corresponding changes in inferred fate distributions.

### Method selection and parameterization

Pseudotime summarizes progression geometry and is ideal for ordering when direction is unknown; it is not a dynamical law and should not be read as physical time or force^1^.

RNA velocity infers short-horizon derivatives from splicing kinetics; scVelo’s dynamical model generalizes to transient regimes (gene-specific rates and latent time). Velocity is most reliable locally and is best used as a directional prior inside probabilistic fate models^2,8^.

OT/Waddington-OT couples distributions across time/conditions to infer probabilistic ancestor-descendant relationships and terminal fate masses. Because couplings are not unique, one must specify biologically motivated priors (growth/death, sampling, cost/regularization) and report sensitivity^3^.

Schrödinger bridges connect entropic OT to stochastic processes on paths, enabling interpolation between marginals with explicit drift-diffusion structure; they can disentangle directed trends from stochastic spread given adequate priors, time stamps or perturbational anchors^4^.

## Results

### What each method commits to (assumptions, strengths, pitfalls)

Pseudotime — Assumptions: manifold continuity and smooth progression; Strengths: geometry/ordering; Pitfalls: misinterpreting axis distance as time or force; embedding sensitivity; branch handling differs across tools^1^.

RNA velocity — Assumptions: splicing kinetics model (steady state or dynamical mode), adequate unspliced coverage, gene selection and normalization; Strengths: local orientation, transient program discovery; Pitfalls: preprocessing sensitivity, global streamline over-interpretation^2,8^.

OT/WOT — Assumptions: cost metric, regularization, growth/death priors, sampling comparability; Strengths: population fate accounting, reprogramming landscapes; Pitfalls: non-uniqueness of couplings without priors; sensitivity to priors; scalability if naively implemented^3^.

Schrödinger bridges — Assumptions: reference process and regularization; Strengths: principled path distribution with drift/diffusion; Pitfalls: identifiability of drift vs diffusion from snapshots; bias from misspecified priors; compute/training stability^4^.

### Case studies

Case 1 — iPSC reprogramming (distributional evolution): Waddington-OT infers couplings between successive days, reveals alternative fates and highlights modulatory TFs/cytokines; control interpretation: interventions that shift coupling mass toward the pluripotent basin increase iPSC probability^3^. (Fig. 3)

**Figure 3.**
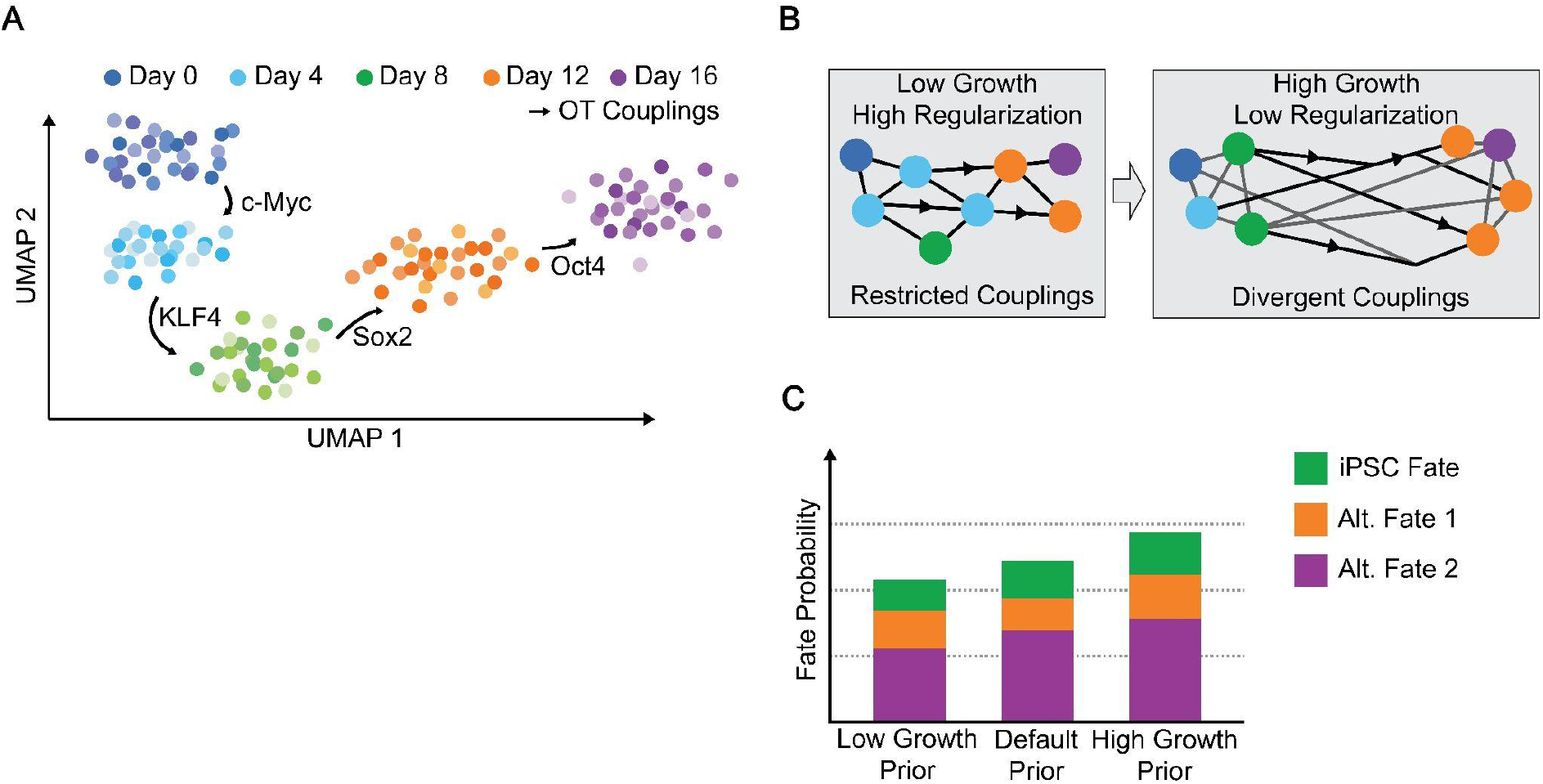
iPSC reprogramming with OT couplings and terminal fate bars with bootstrap intervals. Time-resolved iPSC reprogramming analyzed with Waddington-OT. (A) iPSC reprogramming time course, showing single-cell transcriptional progression during induction with Yamanaka factors: c-Myc, KLF4, Sox2, Oct4. (B) Optimal-transport couplings under varying priors, illustrating how low-growth, high-regularization settings yield more restricted couplings, whereas high-growth, low-regularization settings produce more divergent couplings. (C) Fate probability distributions for iPSC, alternative fate 1, and alternative fate 2, estimated from the inferred couplings.

Case 2 — Pancreas endocrinogenesis (velocity→fate): scVelo’s dynamical mode improves velocity in transients and, with Markov-state fate models (CellRank line), identifies terminal states and fate probabilities; control interpretation: bias transition kernels — reduce dedifferentiation, enhance commitment^8,9^. (Fig. 4)

**Figure 4.**
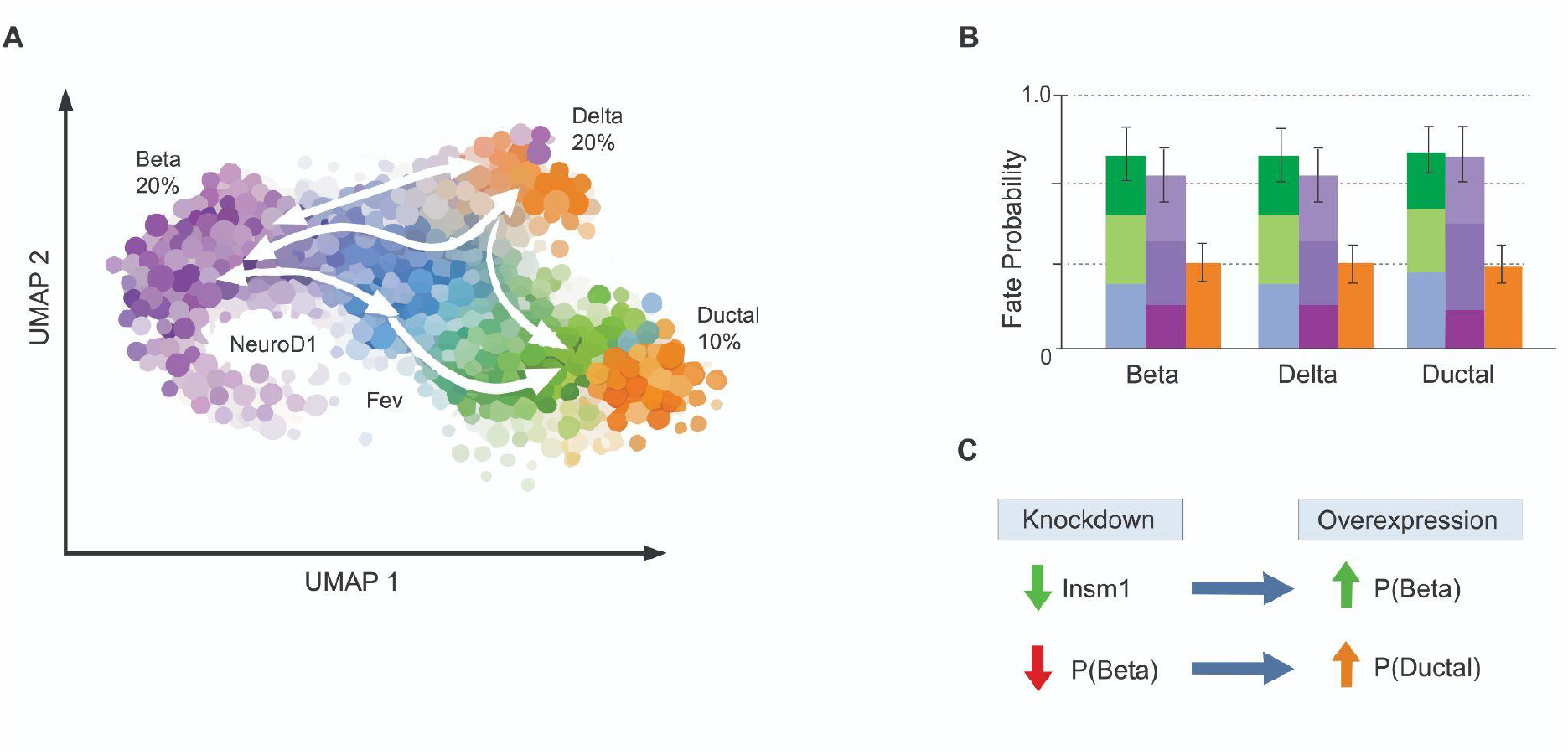
Pancreas differentiation: velocity streamlines; CellRank-style macrostates; fate probabilities. (A) Endocrine differentiation dynamics. RNA-velocity streamlines overlaid on the single-cell embedding highlight local transcriptional flow toward endocrine fates during pancreas development. (B) Optimal-transport couplings under varying priors. Changing growth and regularization parameters alters the coupling structure: low-growth, high-regularization settings produce more restricted couplings, whereas high-growth, low-regularization settings generate more divergent mappings toward β-cell, δ-cell and ductal lineages. (C) Inferred driver genes and perturbation effects. Insm1 emerges as a key regulatory driver. Perturbation analysis shows that Insm1 knockdown reduces the probability of β-cell commitment, while Insm1 overexpression increases β-cell probability and elevates ductal differentiation probabilities, consistent with predicted shifts in transition dynamics.

Case 3 — Hematopoiesis at scale (multiview fate mapping): CellRank 2 unifies velocity, pseudotime, explicit time points and modalities to estimate terminal states and fate probabilities across millions of cells; report compute budgets and bootstrap stability across kernels/priors^9^. (Fig. 5)

**Figure 5.**
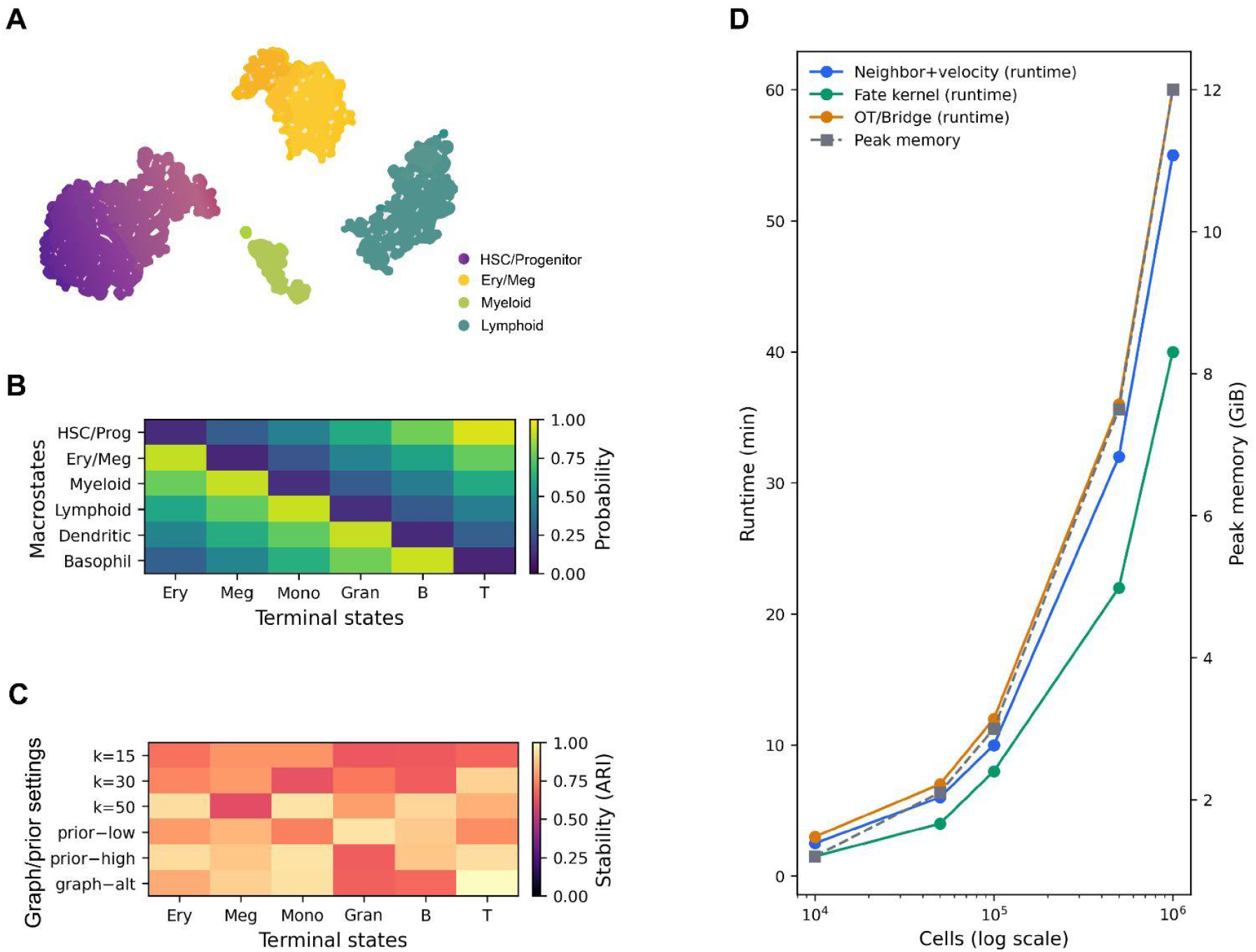
Hematopoiesis at scale: multiview fate mapping and stability analysis. CellRank 2 integrates RNA velocity, pseudotime, explicit time labels and multimodal measurements to estimate terminal states and fate probabilities across large single-cell hematopoietic datasets. (A) Low-dimensional embedding colored by inferred macrostates capturing stable lineage programs. (B) Fate-probability matrix showing commitment likelihoods from progenitor macrostates to terminal states. (C) Bootstrap stability analysis across neighborhood graphs, kernels and prior settings, reporting variability in terminal state assignments and fate probabilities. (D) Computational scaling summary (runtime and peak memory) demonstrating feasibility of million-cell analyses using sparse kernels and batched solvers.

### Applications

Applies to development, regeneration, reprogramming, immune activation/exhaustion, tumor state transitions and organoid maturation—settings where snapshots dominate and interventions are feasible; control goals include increasing desired lineage probability, reducing dedifferentiation or reshaping transient occupancy while preserving viability.

### Reproducibility and data deposition

- Provide complete pipelines (counts→neighbors→embedding→velocity fit→OT/bridge solver→fate estimates) with seeds for stochastic steps.
- Propagate and report uncertainty: subsampling/bootstrapping across preprocessing, neighborhoods and embeddings; uncertainty bands for velocity streamlines; bootstrap intervals for OT couplings and fate probabilities; sensitivity to priors (growth/death/cost).
- Deposit raw and processed data (e.g., AnnData/loom) including kernels, couplings, fate matrices and parameter configurations; link perturbation designs and guide libraries.

### Limitations and optimizations

Identifiability: snapshots alone cannot uniquely separate drift from diffusion nor guarantee unique OT couplings; mitigations: include time/lineage/labeling, impose biologically motivated priors and state uncertainty explicitly.

Stochasticity vs misspecification: diffusion-like spread may reflect genuine heterogeneity or modeling/measurement artifacts; discriminate via controls, replicates, perturbations and ablations.

Scalability: prefer linear/near-linear graph construction, sparse kernels, batched Sinkhorn/bridge solvers, memory-mapped arrays; report runtime and memory transparently.

Integration artifacts: multimodal alignment can distort geometry; validate on held-out structure and propagate alignment uncertainty into fate estimates.

GRN inference limits: multiple networks can explain the same data; propose falsifiable edges and use targeted perturbations to arbitrate among alternatives.

### Outlook

The near-term realistic goal is probabilistic programmability of cell fate: estimate drift/diffusion with uncertainty, nominate levers and test whether perturbations shift terminal distributions as predicted; the long-term opportunity is closed-loop control with online measurements, unifying mechanistic GRN models with distributional estimators and multiview data for safe, robust steering of cell populations.

### Box 1 | Minimal glossary

State space — representation of the cell’s molecular configuration (often latent).

Attractor/basin — stable region toward which dynamics converge (often interpreted as a fate).

Partial observability — we measure y (RNA/ATAC/protein), not the full x.

Vector field — local direction 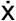 over state space.

Distributional evolution — population dynamics as evolving probability distributions (OT/bridges).

Control input — perturbation (drug/TF/CRISPR) altering dynamics or transition probabilities.

Identifiability — whether distinct models produce indistinguishable observations.

### Box 2 | CME → SDE → Fokker-Planck → quasi-potential (minimal math bridge)

The Chemical Master Equation (CME) models reaction-driven transitions in molecule counts; in mesoscopic limits, a Langevin SDE dx = f(x,u)dt + G(x)dW_t_ approximates the dynamics and the density p(x,t) obeys a Fokker-Planck equation; under small noise, a quasi-potential approximates basin minima and saddles without assuming global gradient dynamics^5,6,7^.

### Box 3 | Observability and feasible control in cells

With y = h(x)+ϵ, exact state reconstruction is impossible; practical objectives target distributional steering under constraints: alter transition kernels or terminal fate distributions (not deterministic paths) while respecting viability, dosage and off-target limits; Perturb-seq makes inputs explicit and measurable, enabling inference of how interventions reshape fate probabilities^10^.

### Box 4 | A 10-step practical checklist (counts → control)

1. QC & normalization
2. Construct neighbor graph
3. Embed (UMAP/PHATE) with stability checks
4. Compute pseudotime
5. Estimate RNA velocity (dynamical mode)
6. Apply OT or bridge models for time series
7. Infer terminal states & fate probabilities (CellRank-like models)
8. Propagate uncertainty (bootstrap/subsample; sensitivity to priors)
9. Nominate regulators (GRN hypotheses) with uncertainty quantification
10. Test via Perturb-seq; evaluate shifts in terminal fate distributions

## Acknowledgements

The authors thank members of the Genomics & Bioinformatics Center at Brigham Young University for helpful discussions that contributed to the development of this framework.

This work was supported by Brigham Young University Life Sciences Faculty Support Grants. The funding source had no role in the conceptual development, writing, or decision to post this manuscript as a preprint.

## AI Assistance Disclosure

Portions of this manuscript were drafted or edited with the assistance of AI-based language tools. The conceptual framework, scientific interpretation, and final revisions were developed and approved by the human authors.

## References

1. Trapnell, C. et al. Pseudotemporal ordering of single cells. Nat. Biotechnol. 32, 381–386 (2014).

2. La Manno, G. et al. RNA velocity of single cells. Nature 560, 494–498 (2018).

3. Schiebinger, G. et al. Optimal-transport analysis of single-cell gene expression identifies developmental trajectories. Cell 176, 928–943.e22 (2019).

4. De Bortoli, V., Thornton, J., Heng, J. & Doucet, A. Diffusion Schrödinger bridge with applications to score-based generative modeling. Adv. Neural Inf. Process. Syst. 34, (2021).

5. Gillespie, D. T. Exact stochastic simulation of coupled chemical reactions. J. Phys. Chem. 81, 2340–2361 (1977).

6. van Kampen, N. G. Stochastic Processes in Physics and Chemistry. 3rd edn (Elsevier, 2007).

7. Ao, P. Potential in stochastic differential equations: novel construction. J. Phys. A 37, L25–L30 (2004).

8. Bergen, V., Lange, M., Peidli, S., Wolf, F. A. & Theis, F. J. Generalizing RNA velocity to transient cell states through dynamical modeling. Nat. Biotechnol. 38, 1408–1414 (2020).

9. Weiler, P., Lange, M., Klein, M., Pe’er, D. & Theis, F. J. CellRank 2: unified fate mapping in multiview single-cell data. Nat. Methods 21, 119–128 (2024).

10. Dixit, A. et al. Perturb-seq: dissecting molecular circuits with scalable single-cell RNA profiling of pooled genetic screens. Cell 167, 1853–1866.e17 (2016).

